# Thermal modulation of skin friction at the fingertip

**DOI:** 10.1101/2022.08.18.504401

**Authors:** Davide Filingeri, Peter Worsley, Dan Bader, Daniel M. Martinez, Konrad Rykaczewski

**Affiliations:** THERMOSENSELAB, Skin Sensing Research Group, School of Health Science, University of Southampton, UK; PRESSURELAB, Skin Sensing Research Group, School of Health Science, University of Southampton, UK; School for Engineering of Matter, Transport and Energy, Arizona State University 501 E Tyler Mall, Tempe, AZ 85287, USA; Julie Ann Wrigley Global Futures Laboratory, Arizona State University, Tempe, AZ, 85287, USA

**Keywords:** Skin Temperature, Friction, Pressure Ulcer, Optical Coherence Tomography, Computational Modeling, Biophysics

## Abstract

Preliminary human studies show that reduced skin temperature minimises the risk of mechanically-induced skin damage. However, the mechanisms by which cooling enhances skin tolerance to pressure and shear remain poorly understood. We hypothesized that skin cooling below thermo-neutral conditions will decrease friction at the skin-material interface. To test our hypothesis, we measured the friction coefficient of a thermally pre-conditioned index finger sliding at a normal load (5N) across a plate maintained at three different temperatures (38, 24, and 16□). To quantify the temperature distribution of the skin tissue, we used 3D surface scanning and Optical Coherence Tomography to develop an anatomically-representative thermal model of the finger. Our data indicated that the sliding finger with thermally affected tissues (up to 8mm depth) experienced significantly (p<0.01) lower frictional forces at 16°C-plate temperature than at the 24°C [-23% (±19% SD)] and 38°C plate interactions [-35% (±11% SD)], respectively. This phenomenon occurred without changes in skin hydration during sliding. Accordingly, our experiments demonstrate thermal modulation of skin friction in the absence of skin-moisture effects. Our complementary experimental and theoretical results provide new insight into thermal modulation of skin friction that can be employed for developing thermal technologies to maintain skin integrity under mechanical loading.

## 1. Introduction

Pressure ulcers (PUs) constitute a localised damage to the skin resulting from prolonged periods of pressure and shear forces, and represent a significant public health challenge, due to their impact on individuals’ quality of life and the associated financial burden on healthcare providers[1]. Indeed, in the United Kingdom alone, the annual cost of treating chronic skin wounds, including PUs, has been estimated to be approximately £8.3 billion[2]. Accordingly, an improved understanding of the fundamental mechanisms underlying the physiological tolerance of human skin to mechanical loading could lead to the development of cost-effective, personalised solutions to promote skin health.

Exposure to prolonged mechanical loading on the skin can arise from several situations, including periods in lying and sitting postures and the prolonged attachment of medical devices. The associated pressure and shear forces acting on the skin induce internal tissue deformations, causing changes to the physiology of skin and sub-dermal tissue. The negative impacts of the deformation include ischemia in the blood vasculature, lymphatic impairment, and direct deformation damage[3]. In addition, the moisture level, and the temperature within and around skin tissues strongly influence skin tolerance to mechanical loading.

For example, elevated humidity at the skin interface reduces the mechanical stiffness and strength of the skin and can increase its friction coefficient[4,5]. In contrast, lower-than-normal temperature reduces skin tissue’s metabolic demands and could increase skin’s physiological tolerance to pressure-induced damage[6]. Indeed, early animal[7] and preliminary human[8,9] studies show that reduced skin temperature minimises the risk of PU formation[10]. While this evidence highlights the potential therapeutic role of skin cooling for maintaining tissue health, the mechanisms by which cooling enhances skin tolerance to pressure, shear, and friction remain poorly understood[11].

Since heating the skin increases its friction at a low normal load (0.5N)[12], we hypothesized that skin cooling below thermo-neutral conditions will decrease friction at the skin-material interface. To test our hypothesis, we measured the friction coefficient of a thermally pre-conditioned human index finger sliding at a normal load (5N) across a plate maintained at three different temperatures. While other body regions (e.g., sacrum, centre of the glut, heel, and neck) are at a greater risk of PU formation[13], the accessibility, ease of movement control, and possibility of localized simulation-designed thermal preconditioning make the finger an ideal site to investigate the relationship between temperature and friction. To quantify the temperature distribution of the skin tissue, we used 3D surface scanning and Optical Coherence Tomography (OCT) to develop an anatomically representative thermal model of the finger. The complementary experimental and theoretical results provide new insight into thermal modulation of skin friction that can be employed for developing thermal technologies to maintain skin integrity.

## 2. Methods

### 2.1 Skin friction experiments

We conducted experiments with air temperature ~24°C and relative humidity ~50% and one healthy male participant (age: 35y; height: 175cm; body mass: 80kg) who was extensively familiarised and trained with the experimental procedures. The participant slid their index finger pad against the surface of a thermally controllable, aluminium friction rig (see **Fig.S1.1**), which was set at either thermo-neutral (24°C), cold (16°C), or warm (38°C) state. The A-to-D data acquisition unit (MC computing, USA) interfaced with the friction rig samples the plate surface temperature, the normal force, and the tangential force at 33Hz. The sliding movement consisted of actively pulling the finger pad from a fixed starting point (i.e., at the top of the plate) to an end point located 100mm away on a straight line (i.e., at the bottom of the plate) at a speed of ~3.3 cm.s^−1^, and was repeated 10 times per plate temperature. The participant was required to maintain a stable 5N normal force during the pulling action, using visual feedback from the digital meter described above. Normal and tangential forces were used to calculate the coefficient of kinetic friction (CoF)[14] and were averaged every 0.33s during each sliding movement. The data was analysed for the independent and interactive effects of plate temperature (3 levels: 24, 16, or 38°C) and time (10 levels: 0 to 0.33s) by means of a 2-way repeated measure ANOVA. We also measured skin temperature (T_sk_) using infrared thermography (IR), and blood flow dynamics using Laser Doppler Flowmetry, and finger pad’ skin conductance that is used to determine local skin hydration level (before and after sliding). Section **S1** of ESM covers the experimental protocol for skin friction experiments in detail.

### 2.2 The geometrical model

We have developed an anatomical model of the distal and middle phalanges of the index finger pressing on a flat surface that combines realistic 3D detail obtained using two imaging techniques[15–19] and geometrically simplified features[20–25]. Specifically, we generated the exterior shape by imaging the finger surface using a white light non-contact handheld scanner (see **Fig.1a-c**). We assumed the epidermis (0.43mm) and dermis (1.1mm) regions to be uniform and conformal to the external surface[18,26] and the fat region to fill the space outside the bones whose shape we obtained from an anatomical database[27]. We created a simplified geometry of the finger contact with the surface based on average of ink imprints and OCT of the air gaps at the interface (see **Fig.1c-e**). The elliptical contact region contains microscale air gaps with a cross-section of a truncated ellipse swept along elliptical segments or circular paths, representing a simplified fingerprint pattern (see **Fig.1c**). The simplified contact has the same percentage of area occupied by air (34%) and skin (66%) as the average of ten ink imprints. Section **S2.1** of ESM covers further imaging, geometrical, and meshing details.

**Figure 1.**
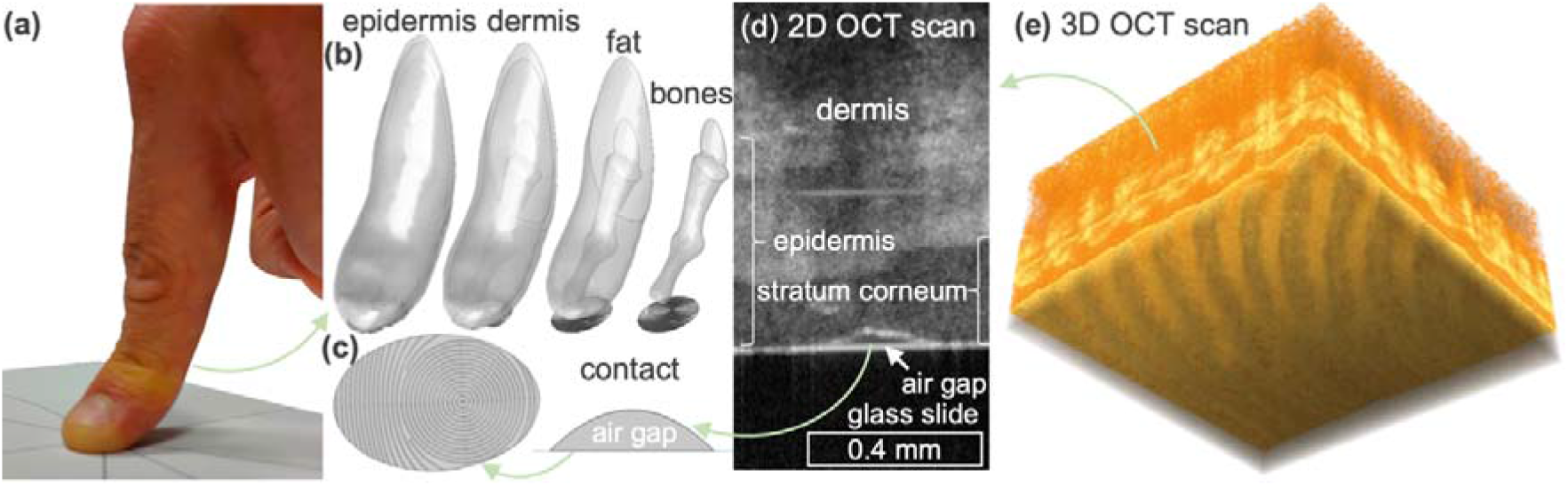
The geometrical model of the index finger in contact with surface: **(a)** photograph of the physical scenario, **(b)** the geometrical model revealing all regions and **(c)** the finger-surface contact, including simplified micro-scale air gap geometry that is based on **(d)** 2D and **(e)** 3D OCT imaging of skin-glass slide contact.

### 2.3 The thermal simulations

We simulated free convection in air and conduction within the finger using the finite element method implemented in Comsol Multiphysics 6.0[27] for the three plate temperatures (). We solved the transient convection first and used the resulting flow and temperature fields as inputs for coupled steady-state convection and transient bioheat transfer simulation. We did not explicitly simulate conduction in the substrate because aluminium experiences a negligible change in surface temperature when in contact with a finger[27]. Instead, we accounted for the substrate with a convection boundary condition at the epidermis-metal interface with the equivalent heat transfer coefficient equal to the inverse of the real-to-projected area adjusted 0.0013 m^2^°CW^−1^ contact resistance[28]. The rest of the simulation setup was based on our prior work[27] and includes skin-to-environment (at 24°C) radiation, isothermal tissue initial condition (at 32°C), and constant boundary condition at the middle and proximal phalanges interface (at 32°C). We benchmarked the simulations against classical semi-infinite analytical model and in more qualitative terms compared against infrared imaging. Section **S2.2** of ESM covers simulation formulation and benchmarking in detail.

## 3. Results

Prior to friction measurements, we thermally pre-conditioned the finger for 60s by pressing it at 5N against the plate with pre-set temperature. Our Laser-Doppler measurements confirmed that the cutaneous blood cell velocity within the pressed finger was substantially reduced to a plate-temperature independent level (see **Fig.S1.3**)[29]. With this reduced blood flow, the external finger temperature within 5-8mm of the substrate changes towards that set for the plate during the 60s period (see simulations in **Fig.2a** and IR in **Fig.S2.4b**). The thermal simulations allow us to quantify the spatial and temporal temperature distribution within the finger during this period. In the early stage of pre-conditioning the internal epidermis temperature is highly heterogenous and follows the spatial pattern of the fingerprint (see **Fig.S2.5b**). The 60s thermal pre-conditioning step reduces the substantial initial temperature gradients in-plane parallel to the substrate (~5°C over 0.2mm) to minor level (under 1°C over 0.2mm, see **Fig.S2.5b**). Consequently, after the pre-conditioning step, we discuss the 3D internal tissue temperature distribution in terms of temperature averaged in-plane parallel to the plate as a function of vertical distance from the plate.

According to the simulation results, the internal finger temperature was impacted up to around 8mm by the 60s contact with the plate. The plot in **Fig.2b** shows that the epidermis (height of 0-0.43mm) is within 1 to 1.5°C of the plate temperature for all the plate settings. From the inner edge of the epidermis layer, the dermis temperature only changed by 1 to 1.5°C when plate temperature was moderately different from the initial skin temperature (i.e., *T*_*p*_ of 24°C and 38°C vs. initial skin temperature of ~32°C). Thus, for *T*_*p*_ of 24°C and 38°C, all of skin tissue (height up to ~1.5mm) is within 2 to 3°C of the plate temperature. At the coldest plate setting of 16°C, the temperature change within the dermis is 3.7°C, so the corresponding skin tissue was within 5°C of the plate temperature. From the inner edge of the dermis (~1.5mm), the temperature of the tissue and bone gradually changed until reaching the initial temperature of 32°C at a depth of ~8mm.

**Figure 2.**
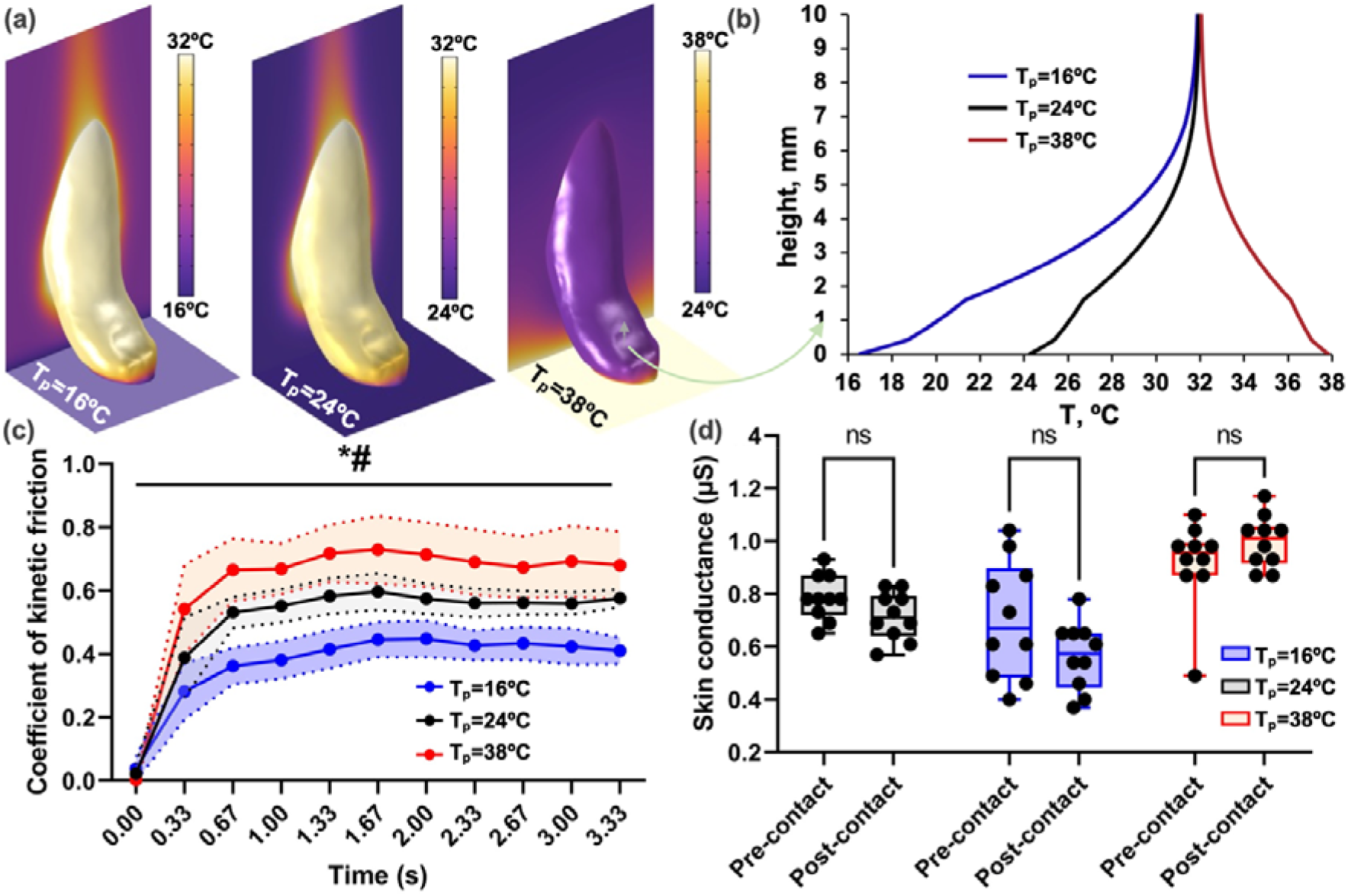
The modelling and experimental results for of 24, 16, and 38°C: **(a)** surface temperature fields, **(b)** temperature of tissue averaged in-plane parallel to the plate as a function of vertical distance from the plate after 60s of pre-conditioning; **(c)** Mean (N=10) ± SD for CoF data recorded during the 10 sliding movements (*denotes main effect of temperature at p<0.05; # denotes main effect of time at p<0.05); **(d)** Box and Whisker plots (Min to Max) for skin conductance data pre- and post-contact (i.e., 10 sliding movements, individual dots) (“ns” denotes non-statistically significant difference at p>0.05).

Our experimental friction measurements indicated that the sliding finger with thermally affected deep tissues (up to 8mm) experienced large and statistically significantly different CoF(s) as a function of plate temperature (temperature main effect: F_(2,18)_=41.01; p<0.0001; **Fig. 2c**). Specifically, CoF(s) during the sliding movement were 0.68 (±0.13 SD), 0.55 (±0.06 SD), 0.40 (±0.07 SD) for the 38°C, 24°C, and 16°C plate temperatures, respectively. By design, normal force during sliding was kept constant at 5N, and this was confirmed experimentally (mean normal force at 38°C: 5.1±0.2N; 24°C: 5.4±0.1N; 16°C: 5.4±0.2N; no temperature main effect: F_(2,18)_=2.15; p=0.145; see **Fig. S1.2**). Hence, frictional forces during sliding movement at 38°C and 16°C plate temperatures (i.e., calculated as CoF x normal force) were 24% (±32% SD) higher and 23% (±19% SD) lower than during the 24°C-plate interaction, respectively. Importantly, we found no differences in skin conductance, and therefore hydration levels, between pre- and post-sliding at each plate temperature (mean difference pre-vs. post-sliding: 0.04µS [95%CI: −0.06, 0.15]; no time main effect: F_(1,9)_=0.79; p=0.397; **Fig. 2d**), which indicated that no sweating-nor condensation-induced moisture appeared at the skin-plate interface during sliding.

## 4. Discussion and Conclusions

Our results demonstrate that skin CoF can be modulated by changing the temperature of the interfacing substrate away from that of the ambient conditions. Our findings expand those of Choi et al.[34] in relation to the effects of pronounced skin cooling (i.e., <23°C) on friction, as we observed frictional forces at 16°C plate temperatures that were 23% (±19% SD) and 35% (±11% SD) lower than during the 24°C- and 38°C-plate interactions, respectively. In contrast to Choi et al.[30], we observed this phenomenon without any change in the skin hydration level during sliding. Accordingly, our experiments demonstrate predominantly thermal modulation of skin friction and not a correlated skin-moisture and temperature induced effect.

Choi et al. have hypothesized that temperature changes the mechanical properties of the tissue (which is generally observed [31,32]) relevant to sliding[30]. Specifically, they observed that increase from 23°C to 42°C did not change the interfacial shear strength and explained the ~50% increase in the CoF by increased contact area of the mechanically-softened tissue[30]. Empirical fitting of their friction data to the Kelvin-Voigt viscoelastic model showed that the temperature increase related to ~143 to 56 MPa viscosity, 15.5 to 10.1 MPa modulus 1, and 5.6 to 5.2 MPa modulus 2 decrease. However, even without changing of the plate temperature from that of the surrounding, sliding changes the contact area of a finger in a complex manner on multiple scales.

Delhaye et al.[33] showed that after ~100ms of sliding, the apparent finger contact area drops by 30-40% (with normal force of 0.5-2N) and shifts its centroid away from the sliding direction. Potentially due to the increased pressure within the reduced apparent area (normal force remaining constant). Liu et al.[34] observed using *in-situ* OCT imaging that sliding of a finger (at 6N or ~20 to 30kPa) increased the microscale substrate contact along a linear scan from 50% to 88% (i.e., sliding shrank the microscale air gaps). We suspect that the highly heterogeneous mechanical properties of the epidermis, dermis, and adipose tissue can impact these processes in complex yet not understood manner. The present study revealed that heat transfer to the deeper tissues is non-linear and temperature dependent. Thus, any thermal induced changes to mechanical properties of skin and fat tissues, and the corresponding contact area, will be complex in nature and requires further multiscale investigation.

The observed thermal modulation of skin friction supports the potential therapeutic benefits of skin cooling regarding minimising friction over vulnerable skin sites. Microclimate conditions at the interface between a loaded skin site at risk of damage (e.g., the sacrum) and a support surface (e.g., a hospital mattress) are commonly associated with ~38°C skin temperatures[35], which is equivalent to the warm-plate condition of the current study. Under this real-life scenario, both our 24°C- and 16°C-plate conditions would represent skin cooling stimuli for a warm, mechanically loaded skin site and they could provide meaningful decreases in friction in a dose-dependent fashion (i.e., −23% at 24°C and −35% at 16°C), resulting from short periods of pre-conditioning (i.e., 60s). This knowledge could translate to inform the design of user-centred medical devices and wearables, including support surfaces and garments delivering cooling to the skin at a level that is both beneficial and comfortable[36]. Indeed, our observed dose-response relationship between skin cooling and friction indicates that friction gains can be obtained at less pronounced levels of cooling (i.e., 24°C instead of 16°C), which are less likely to be associated with cold discomfort[37,38].

Whilst promising, we recognise that our findings apply to the finger and should therefore be interpreted within this small anatomical region. Indeed, the protective effects of cooling to shear-induced damage, as well as its perceptual effects on discomfort, may vary across the human body, due to regional differences in skin biophysics and morphology[39], and in thermoregulatory[40] and perceptual sensitivities[41–43]. Thus, there may be variations in the efficacy of therapeutic cooling associated with the anatomical site. Identifying how the effects of skin cooling vary both physiologically and perceptually over skin sites at greater risk of PU than the finger will therefore be critical to developing “user-centred” thermal-therapy approaches to maintain skin integrity under mechanical stress that are both effective and comfortable. We believe that the methodological approach presented here will provide the experimental and theoretical framework required to establish regional differences in the temperature dependence of skin friction across the body and over skin sites more commonly affected by PUs.

## Supporting information

Electronic Supplementary Material

## Ethics

The experimental part of this research received ethical approval by UoS as part of application ERGOII 73017.

## Data accessibility

Available upon request.

## Authors’ Contributions

DF, KR, DB, PW: conceptualization, DF: experimentation, KR and DMM: mesh generation and thermal modelling, all authors: writing and editing.

## Acknowledgements

KR acknowledges Research Computing at ASU for providing High Performance Computing resources that have contributed to the research results reported within this paper. DF wishes to thank Terence Harvey at the national Centre for Advanced Tribology at Southampton for performing measurements of the surface roughness of the friction rig, and Silvia Caggiari for performing the 3D scanning of the finger.

