## Supplementary material for "Thermal modulation of skin friction at the fingertip": Electronic Supplementary Material

**Electronic Support Material for “Thermal modulation of skin friction at the fingertip”**

**Table of contents**

**S1. Experimental Methods…………………………………………………………………p.2**

***S1.1 Friction rig…………………..…………………………………………………………*p.2**

***S1.2 Protocol of skin friction experiments*………………………………………….………p.2**

**S2. Theoretical Methods……………………………………………………………………p.5**

***S2.1 Generation of the finger model*………………………………………….…………….p.5**

***S2.2 The thermal simulation formulation and benchmarking*……………….…………….p.6**

***S1. EXPERIMENTAL METHODS***

***S1.1 Friction rig***

Figure **S1.1** shows the friction rig, which consists of a force plate and low resistance trackway, incorporating calibrated load cells to record both normal and tangential forces, respectively (FS2050 1500G Load Cell, TE Connectivity, Schaffhausen, Switzerland; LH Series Linear Guides, NSK, Newark, UK; M5 Digital Scale, Dymo, Stamford, CT, US). Force feedback (N) is continually displayed to the participant during interactions using a digital meter interfaced with the load cells (Daisylab, MC computing, USA). A thermally controllable aluminium plate (surface roughness Ra: ~0.3µm) is mounted upon the force plate and incorporates two Peltier elements (ETH-127-10-13 Peltier Module, Adaptive, Kibworth, UK) that allow PID control of surface temperature via closed-loop, custom-written software (Daisylab, MC computing, USA). The centre of the plate holds a 1-mm hole, allowing for the insertion of an optical probe (Moor Instruments, UK) which is situated at the plate interface. The probe is used for the evaluation of skin blood flow via Laser Doppler Flowmetry (LDF), estimated through a flux value (AUs), whilst the finger is statically compressed against the plate.

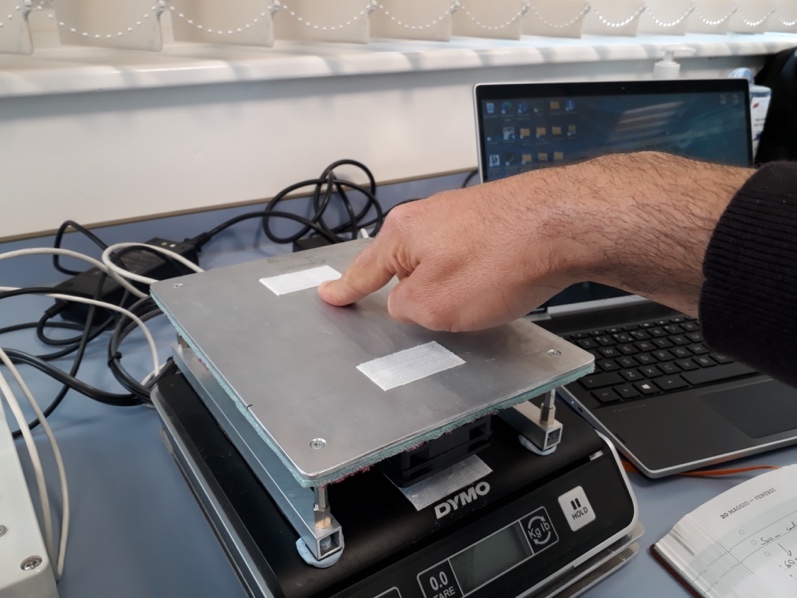

**Fig. S1.1.** Friction rig used in the experiments.

The friction rig is interfaced with an A-to-D data acquisition unit (MC computing, USA), sampling plate surface temperature (°C), as well as normal and tangential forces (N), at 33Hz. Accordingly, we used force data to estimate the kinetic coefficient of friction (CoF). The kinetic CoF can be defined as the resistance between two surfaces moving against each other and was calculated for each sliding movement as the ratio of tangential ($F_{t}$) to normal force ($F_{n}$) according to Amonton’s Law[12]:

$CoF=\frac{F_{t}}{F_{n}}$	**(S.1.1)**

***S1.2 Protocol of skin friction experiments***

During the skin friction experiments, the sliding movement consisted of actively pulling the finger pad from a fixed starting point (i.e. at the top of the plate) to an end point located 100mm away on a straight line (i.e. at the bottom of the plate) at a speed of ~3.3 cm^.^s^-1^. The participant was required to maintain a stable 5N normal force during the pulling action, using visual feedback from the digital meter described above. The consistency in the application of 5N normal force was confirmed by the experimental results (see **Fig. S1.2**)

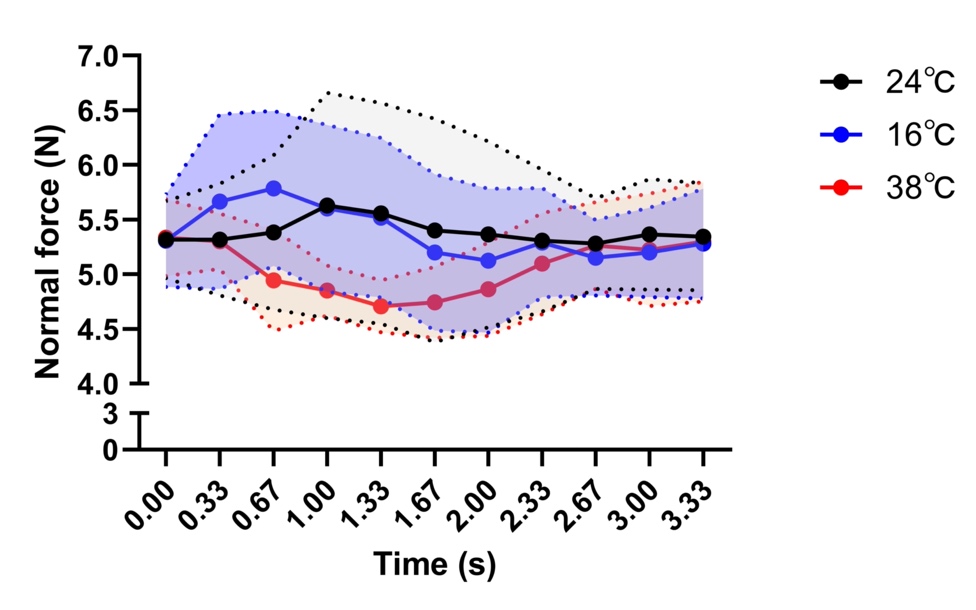

**Fig. S1.2:** Mean (N=10) ± SD for normal force recorded during the 10 sliding movements at each plate temperature (i.e., 24, 16, and 38ºC).

The participant was instructed to first wash and dry their hands. Then a skin conductance (in µS) test was performed on the index finger pad to provide an index of baseline local skin hydration. Second, they were instructed to place their finger at the starting point on the plate and press it down to reach a normal force of 5N. This position was maintained for 60s to allow for adaptation and stabilization of skin temperature in relation to the thermal status of the plate (i.e. set at either 24, 16, or 38ºC). Following this adaptation period, the participant was instructed to pull the finger longitudinally across the plate (as detailed above). The finger was then removed from the plate and placed on the conductance meter to determine post-sliding skin hydration levels. Upon completion of the measurement, the finger was thoroughly dried, and a 1min interval was allowed, before the same sliding sequence was repeated. The participant performed this sliding sequence 10 times at each plate temperature (i.e., 24, 16, or 38ºC).

***S1.3 Skin temperature and blood flow experiments***

For the skin temperature (T_sk_) measurements, infrared images of the finger pad were acquired prior to and immediately after a 60s static contact at 5N normal force against the plate set at either 24, 16, or 38ºC (see **Fig. S2.6**). For the skin blood flow measurements, the finger was first placed on the plate for 60s at a minimal contact force (<1N) to establish baseline skin blood flow without mechanical load. It was then pressed to reach a 5N normal force and monitored for a further 60s. This sequence was repeated at each plate temperature (i.e. 24, 16, or 38ºC).

The T_sk_ and blood flow measurements replicated the contact conditions of the adaptation phase preceding each sliding movement during the skin friction experiments and allowed for an evaluation of the interaction between mechanical load and plate temperature on finger pad skin blood flow (see **Fig. S1.3**).

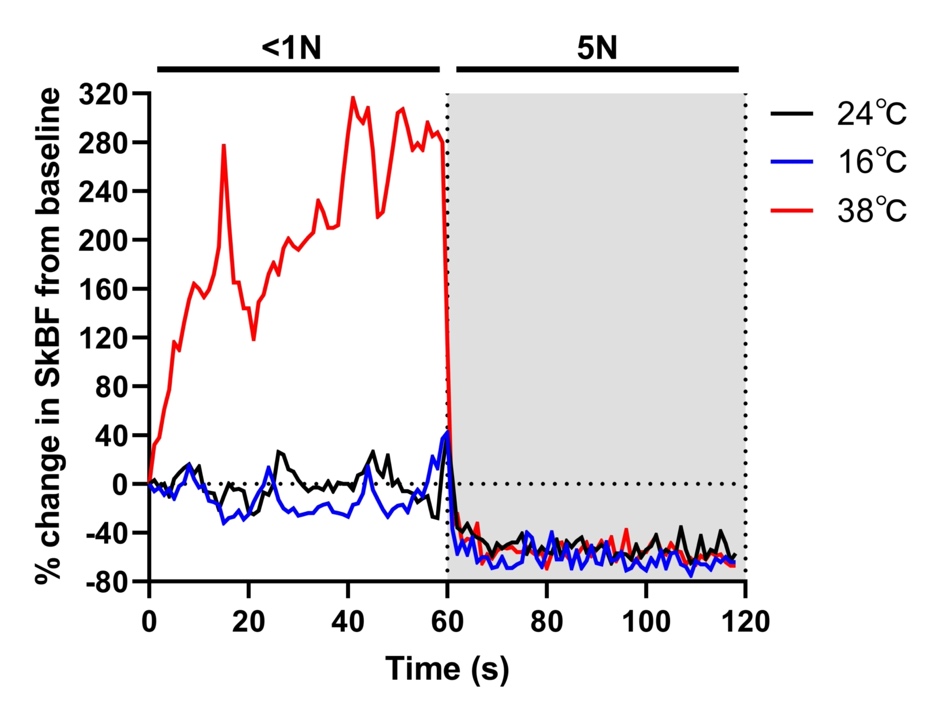

**Fig. S1.3.** Finger skin blood flow data expressed as percentage from baseline values (i.e., arbitrary perfusion units) during minimal mechanical contact (<1N) and during 5N normal force static contact, at each plate temperature (i.e., 24, 16, or 38ºC).

***S2. THEORETICAL METHODS***

***S2.1 Generation of the finger model***

Models of fingers used in previous mechanical or thermal analysis range from highly simplified single or few-layer hemispheres and cylinders [1–6] to more realistic geometries based on tomographic imaging [7–11]. We developed an anatomically representative model of the distal and middle phalanges of the index finger pressing on a flat surface with 5N based on multiscale imaging using multiple instruments. Specifically, the model is based on a combination of three-dimensional macroscale skin surface scanning, anatomical structure database, and microscale optical coherence tomography (OCT; VivoSight Dx OCT System, UK). The geometry consists of bone, fat, dermis, and epidermis regions but neglects finer features such as tendons, muscle, ligaments, microvascular vessels, and joints. We generated the skin surface mesh using a white light non-contact handheld scanner (GoScanner, Creaform) (see **Fig.1a** and **Fig.1b**), which has been demonstrated to have high accuracy and reliability . We processed the scan using Autocad Fusion 360 v2 and Comsol Multiphysics v6.0 to create a watertight mesh with an elliptical contact with the flat surface. We assigned the size of the ellipse (20 mm major axis and 13.4 mm axes with 211 mm^2^ area) based on the analysis of ten ink 5 N imprints of the index finger (see **Fig.1c**). We assumed the epidermis and dermis layers to be conformal with a 0.43 mm (corresponding to the average thickness measured using OCT imaging) and 1.1 mm (based on literature [10,12]) thickness, respectively. We created these meshes using the shell operation in Autocad Fusion. As in our prior work, we obtained the distal and middle phalanx bone shapes from Mathematica V.12.1 anatomical database[13]. We empirically adjusted the angle between the two bones to resemble natural distribution [14] and joined the two bones to create a single innermost region. We assumed the volume in-between the bones and the dermis to be occupied by fat tissue (see **Fig.1b**). We created a simplified geometry of the finger contact with the surface based on the average of OCT measurements of the air gaps at the interface (see **Fig.1c-e**). Specifically, we assumed the air regions to have a cross-section of a truncated ellipse with a 40 µm height (experimental 41$\pm$7 µm with 68% confidence interval) and 160 µm width (experimental 162$\pm$21 µm with 68% confidence interval) and to be spaced by 400 µm. We swept the cross-sections along elliptical segments or circular paths and subtracted the resulting micro-scale features from the epidermis volume (see **Fig.1c**). The final ridge pattern is a simplification of the actual fingerprint. However, it has the same percentage of contact area occupied by air (34%) and (66%) as the average of ten ink imprints. **Fig.S2.1** shows the dimensions of different layers in several views. After mesh refinement study, the final geometry had 4,285,202 elements.

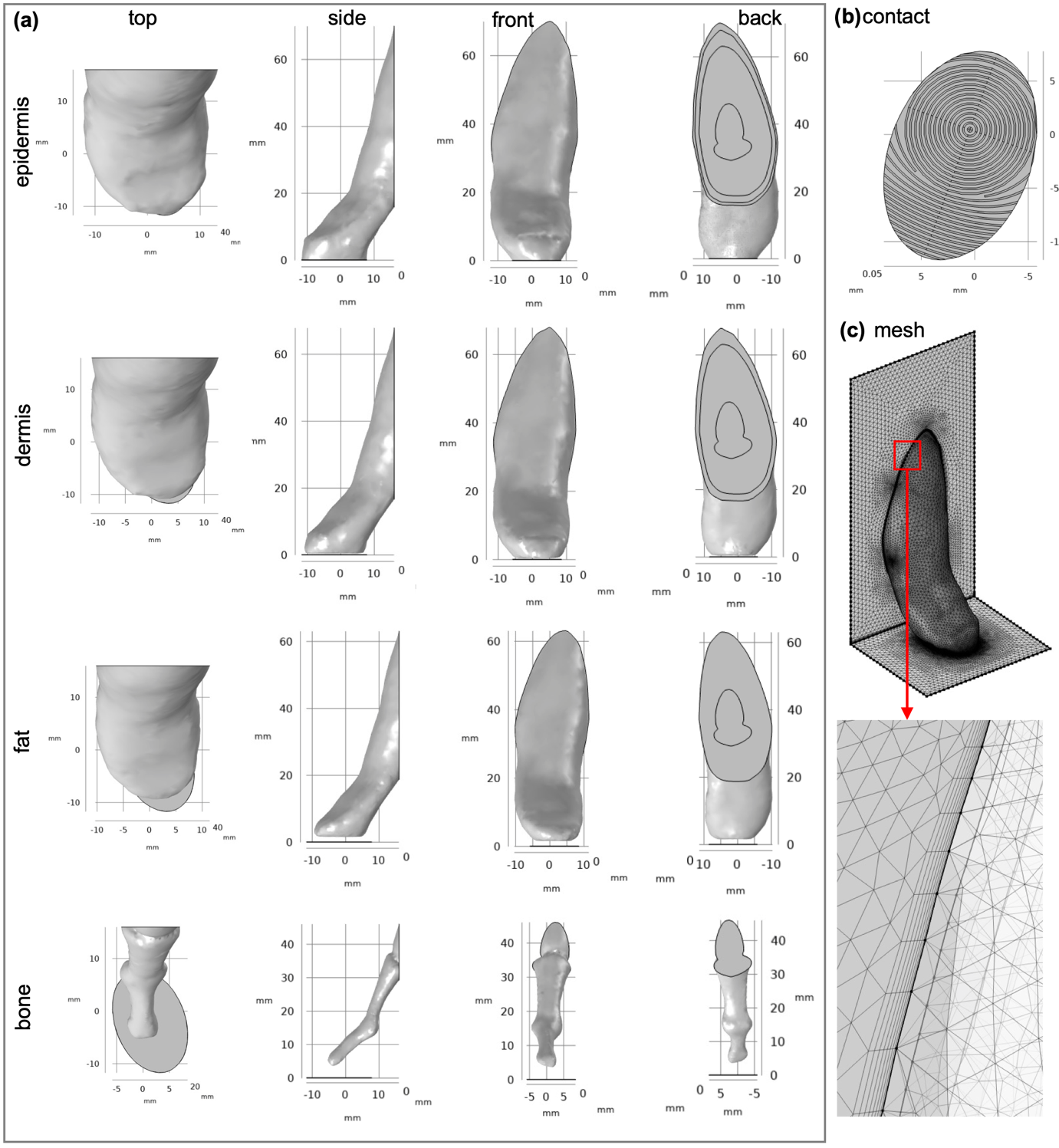

**Figure S2.1.** The geometrical details of **(a)** the anatomical finger model and **(b)** the epidermis-substrate contact; **(c)** the meshed geometry showing boundary layer elements.

***2.2 The thermal simulation formulation and benchmarking***

As in our recent experimentally benchmarked work [13], we simulated natural convection from the finger and the surface and transient bioheat conduction within the finger using finite element analysis (FEA). The simulation had two steps, with the transient convection solved first and used as input for coupled steady-state convection and transient bioheat transfer simulation. For the convection simulation, we assumed the sides and top of the sides of the 50 mm by 50 mm by 100 mm flow region around the finger to be an “Open boundary” condition at the initial air temperature of 24$^{\circ}$C. The plate surface ($T_{p}$ of 16, 24, or 38$^{\circ}$C) was isothermal, and the temperature of the finger vertically varied within 5 mm from the plate temperature to the average of the experimentally observed skin temperature (32$^{\circ}$C—see below in **Fig.S2.4b**). We simulated the buoyancy driven flow by coupling the “heat transfer in fluids” and “laminar flow” interfaces with weakly compressible flow using “non-isothermal flow” multiphysics node in Comsol Multiphysics 6.0. **Fig.S2.2** shows the air temperature and skin heat transfer coefficient distributions for the three plate temperatures 3s into the simulation.

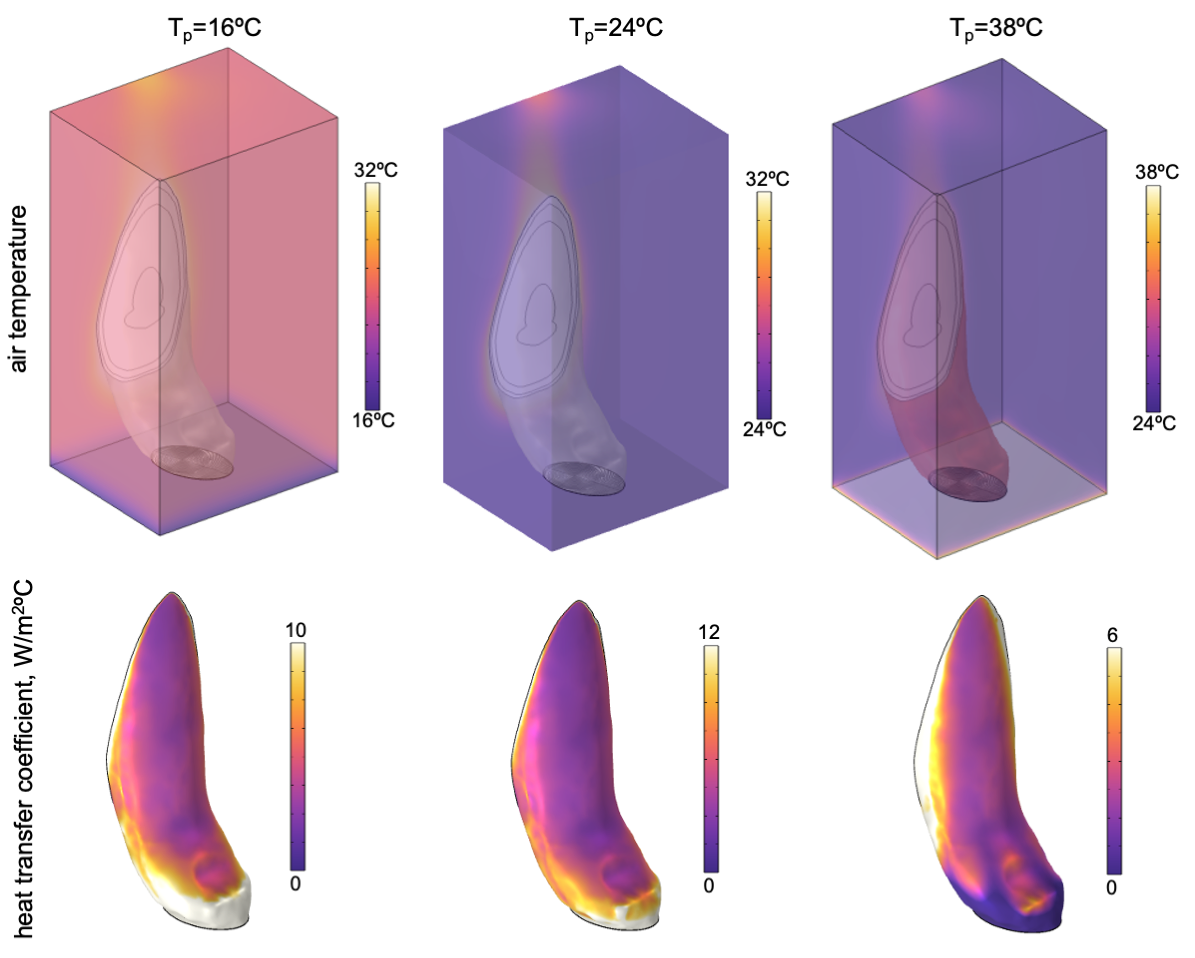

**Figure S2.2.** The steady-state air temperature and heat transfer coefficient distributions for the three plate surface temperatures.

Based on the average heat transfer coefficient of the finger surface ($h_{a}$ defined with respect to air temperature of 24$^{\circ}$C) evolution shown in **Fig.S2.3**, the convective flow achieved steady state within 0.75 to 1 s. The steady-state $h_{a}$was 9.5 Wm^-2^$^{\circ}$C^-1^, 7 Wm^-2^$^{\circ}$C^-1^, and 1.6 Wm^-2^$^{\circ}$C^-1^ for the $T_{p}$ of 16$^{\circ}$C, 24$^{\circ}$C, and 38$^{\circ}$C, respectively. The 7 Wm^-2^$^{\circ}$C^-1^ value for $T_{p}$ of 24$^{\circ}$C is within the 5 to 10 Wm^-2^$^{\circ}$C^-1^ range typical for vertically to horizontally oriented fingers surrounded by stagnant room temperature air (20 to 24$^{\circ}$C) [13].The $h_{a}$ increases to 9.5 Wm^-2^$^{\circ}$C^-1^ when the plate temperature decreases to 16$^{\circ}$C because the temperature of air near the plate is decreased (but the $h_{a}$is defined with respect to the far-field air temperature of 24$^{\circ}$C). Similarly, the finger-surrounding heat transfer is decreased by air heating with the 38$^{\circ}$C plate because of the smaller temperature difference as well as driving buoyancy force. Since any variations to the flow arising from minor finger surface temperature deviation from the assumed form in the convection simulations are likely negligible, we input the steady-state flow conditions into the bioheat transfer simulation.

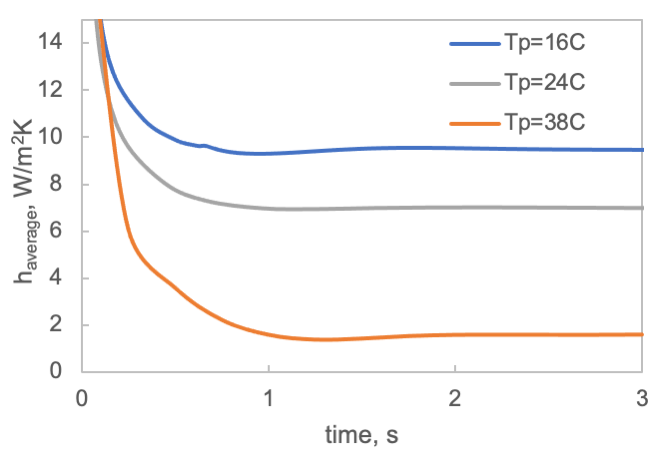

**Figure S2.3.** The evolution of the average convective heat transfer coefficient on the surface of the finger with the three plate surface temperatures specified.

The dermis blood perfusion term ($\omega$) presents the primary unknown for the Penne’s equation-based bioheat transfer simulation. For a subject that “is comfortable as regards to the temperature of the surrounding” [2,15], a blood perfusion rate of 0.0002 to 0.004 s^-1^ [16] and even up to 0.007 s^-1^ [2,15] can occur. Prior research has shown that blood perfusion is decreased by 2 to 8-fold by pressing an index finger against a surface at 5 N [17–19], which we have confirmed using laser-Doppler measurements at the fingertip (see **FigS1.3**). Importantly, the measured velocity increased with temperature in a finger lightly touching a warm surface but was essentially independent of temperature when pressed at 5 N. Since these measurements only provide a relative decrease with finger compression (i.e., the units are arbitrary), we conducted “calibration” simulations of the finger with varied $h_{a}$ and blood perfusion rate. The plot in **Fig.S2.4a** shows that for assumed uniform $h_{a}$ of 5 to 10 Wm^-2^$^{\circ}$C^-1^ (5.6 Wm^-2^$^{\circ}$C^-1^ for the current position without the surface), $\omega$ of 0.004 to 0.006 s^-1^ yields an average skin temperature of 32$^{\circ}$C closely matching infrared imaging (see **Fig.S2.4b**). Accordingly, we used 0.001 s^-1^ as a representative value for the reduced flow. **Table S2.1** lists other thermophysiological properties of the tissue that we obtained from the literature [11,16,20]. We used blood density, specific heat, and temperature of 1050 kgm^-3^, 3770 Jkg^-1^$^{\circ}$C^-1^, and 37 °C, respectively [20].

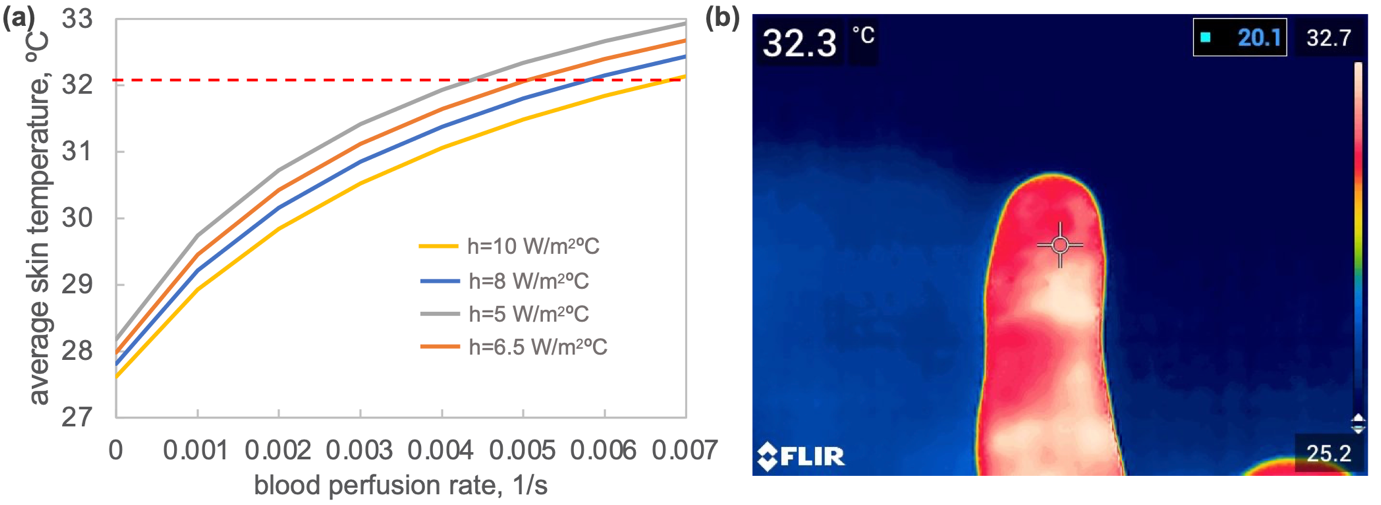

**Fig.S2.4. (a)** the average skin temperature of the finger resulting from bioheat transfer simulations with varied external heat transfer coefficient (assumed uniform) and blood perfusion rate within the dermis and **(b)** illustrative infrared thermal image of the subject's finger prior to experiments demonstrating surface temperature near 32$^{\circ}$C.

**Table S2.1.** Simulated tissue properties [11,16,20]

| Material | Bone | Fat | Dermis | Epidermis |
| --- | --- | --- | --- | --- |
| Thermal Conductivity, $k$, Wm^-1^K^-1^ | 0.32 | 0.21 | 0.44 | 0.23 |
| Density, $\rho$, kgm^-3^ | 1908 | 911 | 1200 | 1200 |
| Specific Heat, $c_{p}$, Jkg^-1^K^-1^ | 1313 | 2348 | 3300 | 3590 |
| Heat generation, $\dot{q}$, Wm^-3^ | 0 | 58 | 368 | 0 |
| Blood perfusion rate, $\omega$, 1/s | 0 | 0.0001 | 0.001 | 0 |

Since our previous simulations showed that the surface temperature of a high thermal conductivity substrate such as aluminium experiences a negligible change when in contact with a finger [13], we did not explicitly simulate conduction in the substrate. Instead, we accounted for it with a convection boundary condition at the epidermis-metal interface with the equivalent heat transfer coefficient equal to the inverse of the for area contact resistance ($R_{c}^{"}$). For aluminum-skin contact with about 25 kPa pressure corresponding to the 5 N force and 211 mm^2^ projected area ($A_{p}$ corresponding to the ellipse area), the area contact resistance is 0.002 m^2^$^{\circ}$CW ^-1^ [21]. Since we explicitly modelled the real skin area that is in contact with the substrate (i.e., the compressed ridges with ${A_{r}=0.66A}_{p}$), we modified the per area contact resistance ($R_{c-r}^{"}$) to provide the same absolute value as for the projected area, namely;

$R_{c}=R_{c}^{"}/A_{p}=R_{c-r}^{"}/A_{r}$ so $R_{c-r}^{"}=R_{c}^{"}A_{r}/A_{p}=0.66 R_{c}^{"}=$ 0.0013 m^2^$^{\circ}$CW ^-1^

The rest of the simulation setup was based on our prior work [13] and includes skin-to-environment (at 24$^{\circ}$C) radiation, isothermal tissue initial condition (at 32$^{\circ}$C), conduction-only treatment of the air gaps, and constant boundary condition at the middle and proximal phalanges interface (at 32$^{\circ}$C).

To benchmark our model, we compared the simulated average epidermis temperature at $z=$ 100 µm above the substrate interface to the classical semi-infinite analytical model with a convective boundary condition ($h=1/R_{c-r}^{"}$) [22]:

$T_{epidermis}\left( z,t \right)=T_{i}+(T_{p}-T_{i})\left\{ erfc\left( \frac{z}{2\sqrt{\alpha t}} \right)-\left( e^{\frac{hz}{k}+\frac{h^{2}\alpha t}{k^{2}}} \right)\left( erfc\left( \frac{z}{2\sqrt{\alpha t}}+\frac{h\sqrt{\alpha t}}{k} \right) \right) \right\}$	**(S.2.1)**

Where $T_{i}$ is the initial temperature of 32$^{\circ}$C, while $\alpha$ and $k$ are the thermal diffusivity (5.3*10^-8^ m^2^s^-1^) and conductivity of the epidermis (see **Table S2.1**). The plot in **Fig.S2.5a** shows that despite the geometrical complexity of the finger model, the highly simplified semi-infinite epidermis model matches closely with simulation results. The minor discrepancies between the simulated and analytical temperatures likely stem from the finger-ridge geometry following temperature heterogeneities in epidermis temperature at $z=$ 100 µm surface. **Fig.S2.5b** shows that these differences can be as high as 5$^{\circ}$C shortly after the contact is established (0.75 s temperature maps) but decrease to 1 to 1.5$^{\circ}$C after 60s.

**Fig.S.2.6** shows a more qualitative benchmarking approach that compares infrared images of the fingertips's palmar surface after sliding and removing the finger from the plate to similarly oriented simulation results after 60s of the contact. In order to corroborate between the two data sets, we had to create a custom colormap for Comsol Multiphysics based on the FLIR Rainbow color map (the match is close but not exact). In the case of the 38$^{\circ}$C plate temperature, the simulated and imaged temperature distributions closely resemble each other. In the other two cases, the match is not as close. However, the main features, such as the ~5 mm transition between the directly cooled and uncooled skin surface areas, are evident. The differences between the simulated and imaged temperature distributions are not surprising since the latter were taken after the finger slid and was removed from the surface (i.e., the finger surface had the opportunity to either cool or heat for a period before the image was taken).

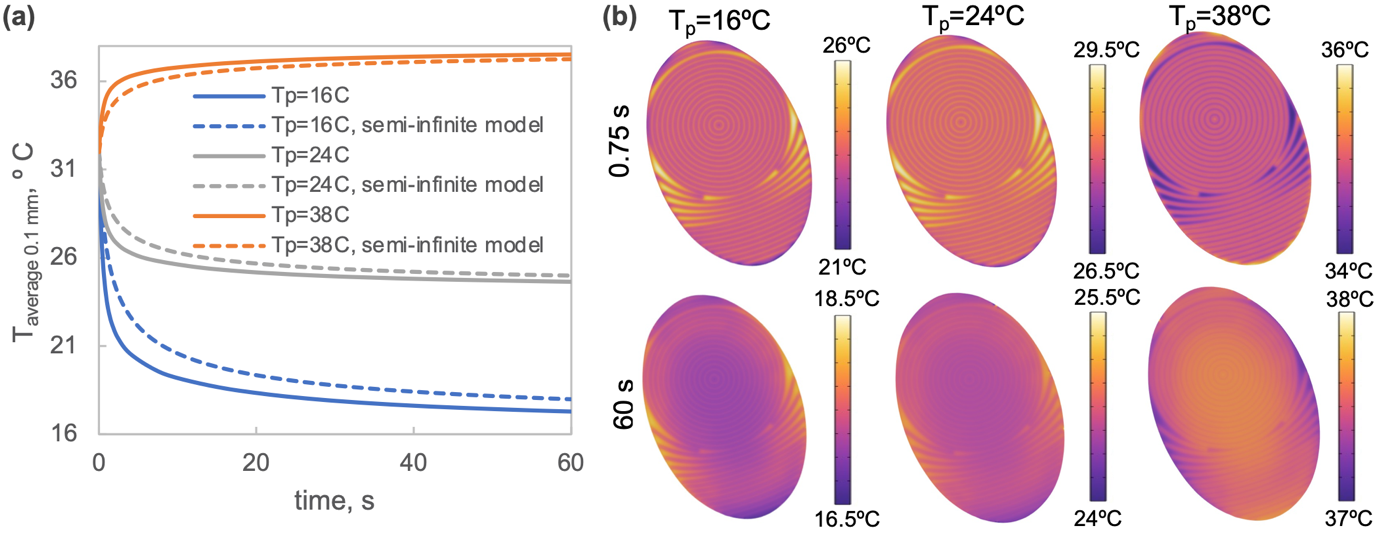

**Fig.S2.5. (a)** plot comparing simulated average skin temperature of epidermis 100 µm above the substrate with corresponding semi-infinite model and **(b)** the temperature fields for the epidermis area 100 µm above the substrate 0.75 s and 60 s after contact with plate at the three different temperatures is established.

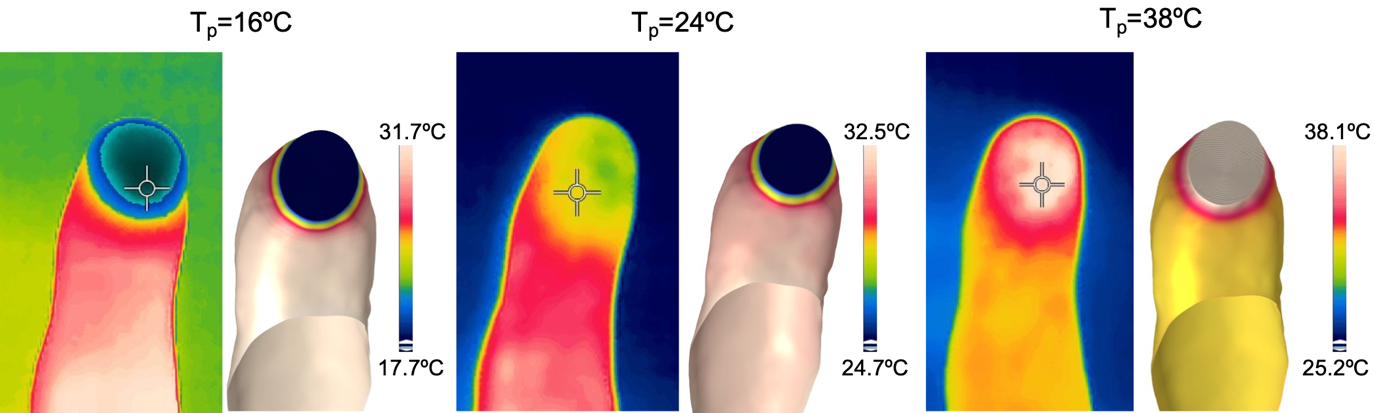

**Fig.S2.6.** Comparison between infrared images of the palmar side of the index finger taken shortly after the sliding experiment and of simulated skin surface temperature after 60s of cooling/heating (but in contact with the plate which was hidden in the image). The colormaps and scales in the experimental and simulated temperature maps were synchronized.
